# RecGraph: adding recombinations to sequence-to-graph alignments

**DOI:** 10.1101/2022.10.27.513962

**Authors:** Jorge Avila, Paola Bonizzoni, Simone Ciccolella, Gianluca Della Vedova, Luca Denti, Davide Monti, Yuri Pirola, Francesco Porto

## Abstract

The transition towards graph pangenomes is posing several new challenging questions, most notably how to extend the classical notion of read alignment from a sequence-to-sequence to a sequence-to-graph setting. Especially on variation graphs, where paths corresponding to individual genomes are labeled, notions of alignments that are strongly inspired by the classical ones are usually able to capture only variations that can be expressed by mismatches or gaps, such as SNPs or short insertions and deletions.

On the other hand the recent investigation of pangenomes at bacterial scale (Colquhoun et al, 2021) shows that most tools are tailored for human pangenomes and are not suited to bacteria which exhibit, among other characteristics, a larger variability. Such variability leads to the need for incorporating a greater flexibility when computing an alignment.

In this paper, we extend the usual notion of sequence-to-graph alignment by including recombinations among the variations that explicitly represented and evaluated in an alignment. From a computational modeling point of view, a recombination corresponds to identifying a new path of the variation graph which is a mosaic of two different paths, possibly joined by a new arc.

We provide a dynamic programming algorithm for computing an optimal alignment that allows recombinations with an affine penalty. We have implemented our approach with the tool RecGraph and we have analyzed its accuracy over some over some bacterial pangenome graphs.

## 1 Introduction

Sequence-to-graph alignment is one of most important computational problems in pangenomics [4]. Still, the field does not have a shared formalization of what is a “good” pangenome graph representing a set of genome sequences [2]. Indeed, instead of optimizing a specific objective function, the currently used approaches are based on heuristics that can scale to manage from population-scale human genomes. The sequence-to-graph alignment problem shows similar issues: the theoretical foundations of the problem are understudied, while the focus is on heuristics that are able to quickly compute a high-quality alignment.

A seminal paper on the foundations of sequence-to-graph alignment is [12], where a notion of alignment of a sequence against a directed acyclic graph has been introduced. The resulting computational problem has been called partial order alignment (POA) with the goal of providing a practical solution to the multiple sequence alignment (MSA) problem, by iteratively adding sequences to the graph with a dynamic programming approach that extends the Needleman-Wunsch algorithm [17] to directed acyclic graphs. In fact, a sequence-to-graph alignment can be used to point out how to change the graph so that it is able to also express the sequence. While this paper predates computational pangenomics, POA has recently gained a renewed interest thanks to its ability to model and attack the sequence-to-graph alignment problem. Some practical improvements have recently appeared, most notably abPOA [9] and Gwfa [25] which incorporate recent advances in dynamic programming alignments and SIMD instructions.

If the pangenome graph has cycles, the alignment problem becomes more complex and some formulations are NP-complete [11]. In that formulation, mismatches are represented as changes in the graph and/or in the sequence, and changes in the graph are needed for the NP-completeness. If changes are allowed only in the sequence, there is a *O*(|*V* | + *q*|*E*|)-time algorithm, where *q* is the size of the sequence and each vertex in *V* is labeled by a single character. Aligning against general graphs can be reduced to finding a shortest path in an alignment graph that is built from the pangenome graph and the sequence to align. Even outside computational pangenomics, finding approximate pattern matching in a graph has attracted interests, starting from the seminal papers [1, 16] and going on with other important complexity results and algorithms for different variants [22, 5]. More recently, the field has found new unsolved problems and important contributions [19, 21, 20].

The need for efficient tools for the sequence-to-graph problem has focused on decreasing the memory usage, employing very efficient data structures such as the Graph Positional BWT [21], and on heuristics that are able to scale to genome-wide graphs [19, 10]. The distinction between *variation graphs* and *sequence graphs* seems to be crucial. The main difference is that variation graphs consider haplotype information represented as distinguished paths [21], while *sequence graphs* do not distinguish paths [9]

On population-scale human pangenome graphs, the *O*(|*V* | + *q*|*E*|) time complexity of [11], limits the practical usefulness of that approach, which justifies the fact that heuristics are much more common on those data. Nevertheless, pangenomics is becoming relevant in the analysis of bacterial and viral species since the degree of variability in these species is even higher [8]; while two human genomes are over 99% alignable, two bacteria of the same species might be 50% alignable [3].

Bacteria frequently import genes, or fragments of them, in place of existing homologous genetic material in their genome, a process that was first identified by the observation of mosaic genes at loci encoding antigens or antibiotic resistance [6, 24], these exchanges of material are known as homologous recombination (HG) and horizontal gene transfer (HGT). There have been significant efforts to understand and study bacterial pangenomes [8]. Some tools are specialized in gene regions [18], while others are focused in intergenic regions, due to the evidence that variation in these regions in bacteria can directly influence phenotypes [23]. Pangenome graph structures have been proposed recently [3]. Nevertheless, a recombination from the alignment point of view has not been yet defined. As such, a main challenge in pangenomics is the construction of a graph that gives evidence of the mosaicism present in a species.

In this paper we explore a first extension of the notion of sequence-to-graph alignment that exploits the fact that a pangenome graph represents a set of related individual or species. More precisely, we introduce the possibility that the sequence has been extracted from a genome that is the result of a recombination between two of the genomes represented in the graph. From a combinatorial point of view, the conceptual approach is similar to the one followed by POA or abPOA, where the result of an alignment is interpreted as a sequence of graph modifications that allow the updated graph to be fully consistent with the read. In our case, we explicit model a recombination as the addition of a new arc and we describe a dynamic programming approach to compute an optimal sequence-to-graph alignment that allows a set of recombinations. We have implemented our approach to compute optimal sequence-to-graph alignments that allow a recombination, with an experimental analysis on a bacterial graph pangenome.

Even though a dynamic programming approach is unlikely to scale to genome-wide graphs, that is not a great limitation of our approach. In fact, almost all current alignment tools are based on a seed-and-extend strategy, where some exact (errorless) matches between the sequence and the graphs are first computed, those matches are chained, and finally the gaps within matches are filled in with a precise, but not necessarily fast, approach. Our main focus has been on this final task, where scalability to huge instances is not a requirement. Finally, an experimentally study of RecGraph over a bacterial pangenome shows that RecGraph is effective in assessing the quality of the alignment of reads from paths that are not represented in the pangenome graph and may be explained as a mosaic effect of path recombinations.

## 2 Preliminaries

Given an alphabet Σ, and *s* an *n*-long sequence (or string) *s* = *s*[1] … *s*[*n*] over Σ, the substring *s*[*i* : *j*] denotes the portion of *s* from the *i*-th character to the *j*-th character, that is *s*[*i* : *j*] = *s*[*i*] … *s*[*j*]. The *k*-long prefix of *s*, that is the string *s*[1 : *k*] is denoted as *s*[: *k*], while the *k*-long suffix of *s*, that is the string *s*[*n* − *k* + 1 : *n*] is denoted as *s*[*n* − *k* + 1]. In this paper, we consider the notion of a variation graph that is a directed acyclic vertex-labeled graph, whose paths correspond to the genome sequences that we want to encode [2, 10]. We refer the reader to [7] for the terminology on graphs.

### Definition 1

(Variation graph). *A* variation graph *G* = ⟨*V, A, W, λ*⟩ *is a directed graph whose vertices are labeled by nonempty strings, with λ* : *V* → Σ^+^ *being the labelling function, and where A denotes the set of arcs and W denotes a nonempty set of distinguished paths, called* walks.

In the following we assume that the DAG has a source node *s*_*i*_ and a sink node *s*_*e*_ and thus all paths in *W* start in *s*_*i*_ and end in *s*_*e*_. Observe that the source and the sink may be labelled by the empty string since they may be added at the beginning and end of each walk in the graph. Given a variation graph *G* and one of its paths *w*, the *path label* of *w* = ⟨*w*_1_, …, *w*_*k*_⟩ is the concatenation of the labels of the nodes in the path *w*, that is the string *λ*(*w*_1_)*λ*(*w*_2_) … *λ*(*w*_*k*_). With a slight abuse of language, we use *λ*(*w*) to denote the path label of the path *w*. Moreover, we focus on acyclic variation graphs, where no path starting and ending in the same vertex is allowed — expect for paths that contain no arc. Notice that this constraint is stronger than asking all walks in *W* to be acyclic. To simplify the presentation, we will only consider canonical variation graph where vertices are character labeled instead of sequence labeled. It is possible to prove that considering only those graphs is not restrictive. In fact, some software tools available (e.g. abPOA [9]) convert the input graph into a canonical graph (see Figure 4).

**Figure 1:**
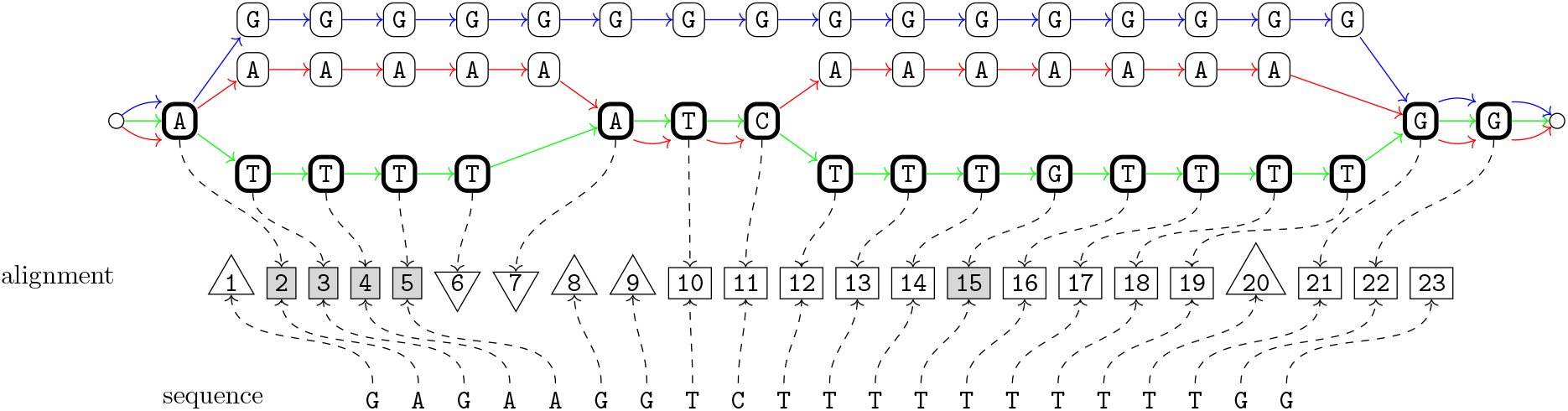
Example of path-preserving alignment of a sequence against a variation graph. The figure consists of three parts: the variation graph is above, the sequence is below, and a representation of the alignment is in the middle. The alignment consists of 23 pairs, where the numbers refer to the pairs and the elements in each pair are represented via a dotted arc that connects the character of the path and/or the character of the sequence that form the pair. This connection is inspired by the notion of threading scheme of [14]. In the representation of the alignment, squares with a white background represent a match, squares with a grey background represent a mismatch. Triangles represent gaps, where a vertex points towards the graph or the sequence to show where an indel is inserted.

**Figure 2:**
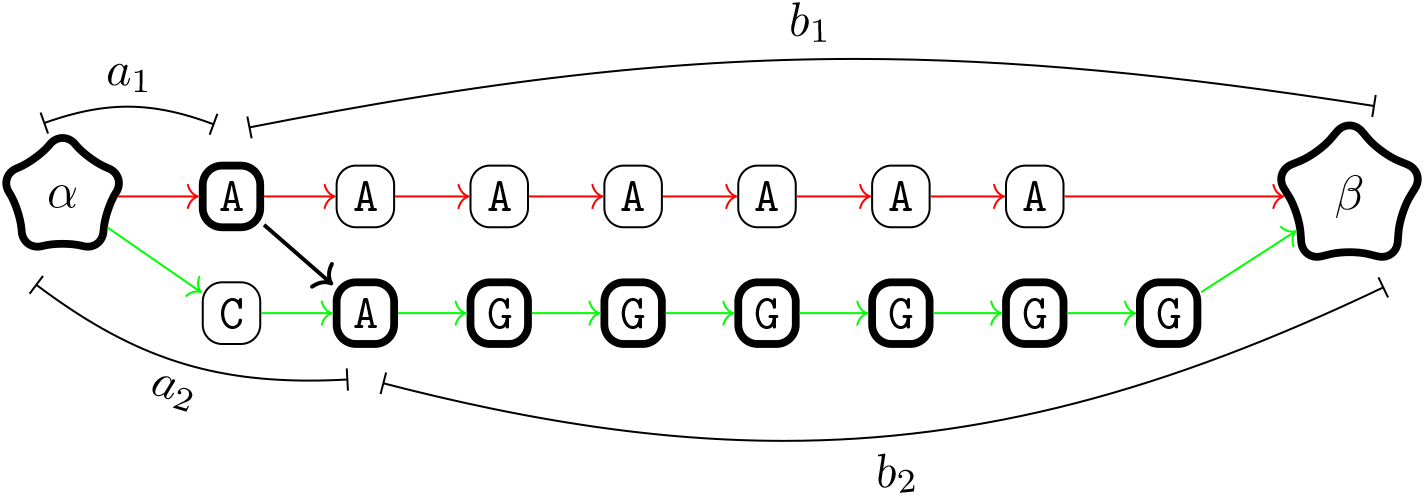
Displacement of the recombination in Figure 3. Only the portion of the graph between the *α* and the *β* nodes are represented. The recombination is represented by the thick black arc connecting two vertices with gray background. The subpaths *a*_1_, *a*_2_, *b*_1_, and *b*_2_ are those of Definition 8. Observe that a recombination may also use an existing arc in the graph in which case the arc represents a switch between two distinct colored paths.

**Figure 3:**
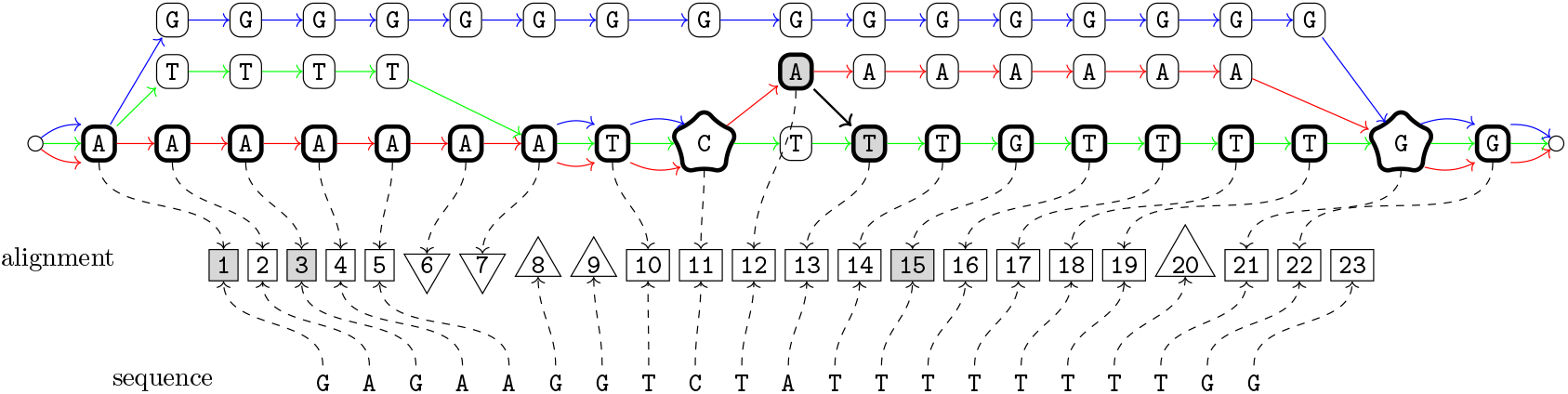
Example of path-preserving alignment of a sequence against a variation graph with a recombination. The recombination is represented by the thick black arc connecting two vertices with gray background. The nodes *α* and *β* are represented by stars. The nodes of the walk *w* have thick edges.

**Figure 4:**
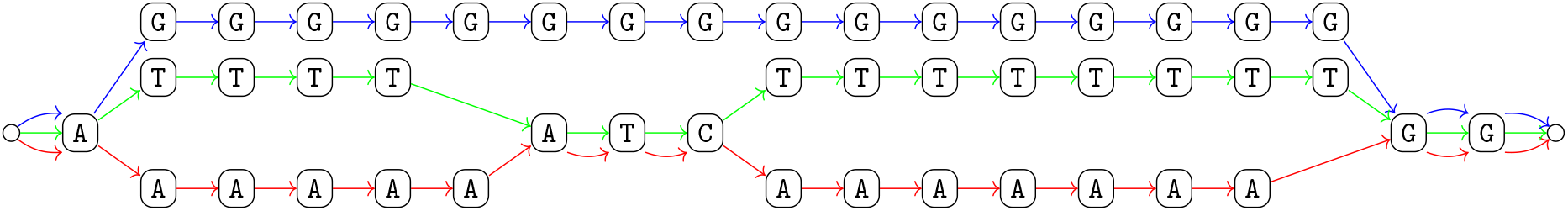
Example of canonical variation graph with three paths (one for each color).

**Figure 5:**
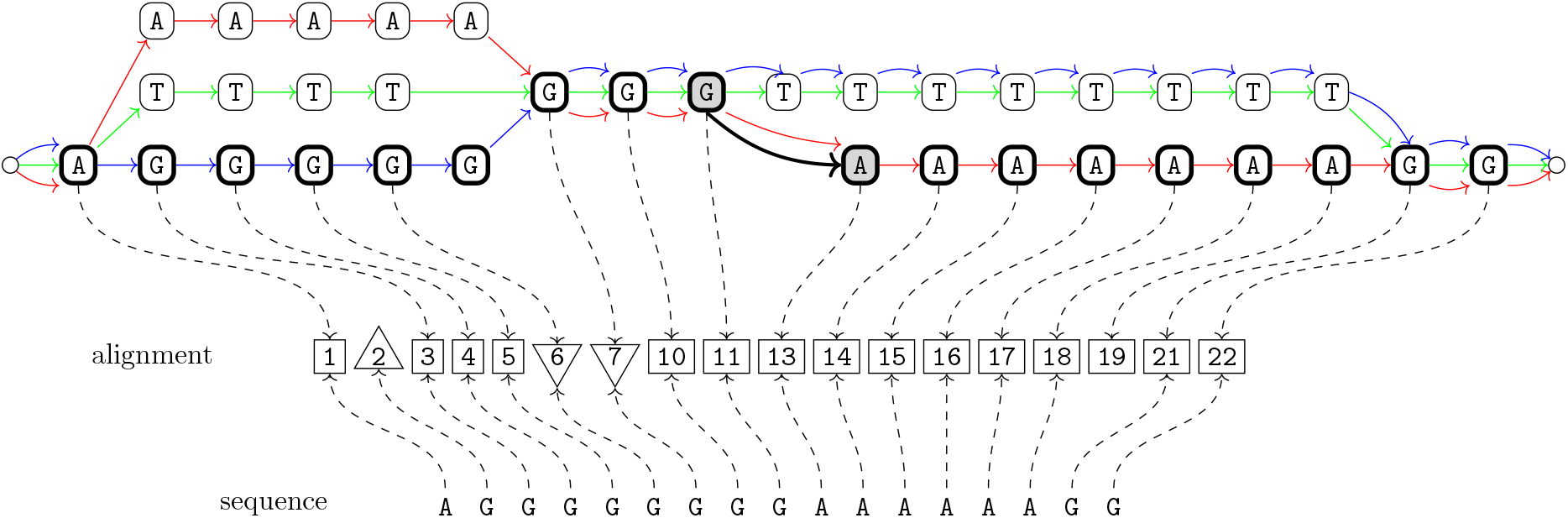
Example of path-preserving alignment of a sequence against a variation graph with a recombination. The recombination is represented by the thick black arc connecting two vertices with gray background. This example shows a recombination arc that is parallel to an already existing arc.

### Definition 2

(Canonical variation graph). *Let G* = ⟨*V, A, W, λ*⟩ *be a variation graph. Then G is* canonical *if all its vertices are labeled by single characters, that is the labeling function is λ*^′^ : *V* → Σ.

In the following, we will consider only canonical variation graphs. Lemma 1 (in the Appendix) shows that this is not restrictive. In the literature, the notion of *sequence graphs* is sometimes used: this corresponds to a variation graph where *W* consists of all possible source-to-sink walks [2].

## 3 Sequence-to-graph Alignment

In the following we define an alignment of a string against a graph as a sequence of pairs of positions of a walk of the graph and of the string. Then a cost of the alignment and the path-constrained condition for a variation-graph will be specified by first defining the notion of graph gap and string gap. Each pair can have an empty position (in the case of a gap), but they cannot both be empty. We denote an empty position with −. Observe that our definition of alignment aims to exploit the structure of the graph mainly when we explore the notion of recombination. Indeed, by viewing the alignment as a sequence of pairs we are able to associate a penalty to an event of recombination in the alignment based on the ordered paired sequences of the two walks involved.

### Definition 3

(Alignment of a string against a variation graph). *Let G* = ⟨*V, A, W, λ*⟩ *be a canonical variation graph, and let s be a string of length l. Then, an alignment of s to G consists of (1) a walk* ⟨*w*_1_, … *w*_*q*_ ⟩ ∈ *W, (2) a sequence* ⟨(*x*_*i*_, *y*_*i*_)⟩ *of ordered pairs where each x*_*i*_ ∈ [1, *q*] ∪ {−} *and each y*_*i*_ ∈ [1, *l*] ∪ {−} *such that:*

1. *if i < j and both x*_*i*_, *x*_*j*_ *are different from* −, *then x*_*i*_ *< x*_*j*_;
2. *if i < j and both y*_*i*_, *y*_*j*_ *are different from* −, *then y*_*i*_ *< y*_*j*_;
3. *each pair has at least an element that is not* −;
4. *for each j* ∈ [1, *q*] *there is an i such that x*_*i*_ = *j, and for each j* ∈ [1, *l*] *there is an i such that y*_*i*_ = *j*,

Informally, conditions (1) and (2) of Definition 3 implies respectively that the alignment is a consistent left-to-right comparison of the string and the graph, while condition (3) corresponds to the usual requirement that no column of an alignment contains only indels. Notice that Definition 3 describes a *global* alignment, where the entire sequence and the entire walk are aligned. In the case of read mappers, it is more common to consider semi-global alignments, where the entire sequence is aligned against a portion of a walk. Again, to simplify the presentation, our definitions will be on global alignments, but the extension to semi-global alignments is quite straightforward. Usually, the presence of indels is penalized with an affine gap penalty. This requires the introduction of the notion of gap and of its length.

### Definition 4

(Gap). *Given an alignment* ⟨(*x*_*i*_, *y*_*i*_)⟩ *(see Definition 3) with z ordered pairs, a* graph gap *consists of a maximal interval* [*b, e*] ⊆ [1, *z*] *such that all x*_*i*_ *with b* ≤ *i* ≤ *e are equal to* −, *while a* string gap *consists of a maximal interval* [*b, e*] ⊆ [1, *z*] *such that all y*_*i*_ *with b* ≤ *i* ≤ *e are equal to* −. *The* length *of such a gap is equal to e* − *b* + 1, *and will be denoted by l*(*b, e*).

The value of an alignment depends also on a score matrix *d* that assigns a value to each pair of characters, and a penalty *g*(*l*) for each gap long *l*. In practice, we will consider only affine gap penalties.

### Definition 5

(Value of an alignment). *Let s be a sequence, let G* = ⟨*V, A, W, λ*⟩ *be a variation graph, and let w*, ⟨(*x*_*i*_, *y*_*i*_)⟩ *be an alignment of s and G. Assume that* ⟨(*x*_*i*_, *y*_*i*_)⟩ *has z ordered pairs and k gaps* [*b*_1_, *e*_1_], …, [*b*_*k*_, *e*_*k*_], *and let B* = {*j* ∈ [1, *z*] : *j* ∉ [*b*_*i*_, *e*_*i*_] *for any i* ∈ [1, *k*]}. *Then the* value *of the alignment is the sum*

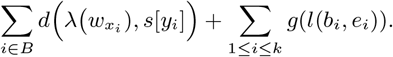

In Definition 5 *w*_*k*_ is the *k*-th vertex of the walk *w* of the alignment, see Definition 3. The first component of the value is the sum of the values of all columns that are not part of a gap, while the second part is the sum of all gap penalties.

### 3.1 Alignments with recombinations

In this section we extend the notion of path-preserving alignment to also allow recombinations. The main idea is that the result of a recombination is a mosaic of two of the walks of the variation graph *G*, where such a mosaic is not a walk of *G*. First we introduce the notion of alignment of a string against a walk; this definition is a variant of Definition 3 where the graph is a single path which is not restricted to start and end in the source and the sink.

#### Definition 6

(Alignment of a string against a walk). *Let G be a character-labeled dag consisting of a single walk, let v*_1_ *and v*_2_ *be respectively the source and the sink of G, and let s be a string with length l. Then an alignment of s to the G consists of (1) a sequence* ⟨(*x*_*i*_, *y*_*i*_)⟩ *of ordered pairs where each x*_*i*_ ∈ [1, *q*] ∪ {−} *and each y*_*i*_ ∈ [1, *l*] ∪ {−} *such that:*

1. *if i < j and both x*_*i*_, *x*_*j*_ *are different from* −, *then x*_*i*_ *< x*_*j*_;
2. *if i < j and both y*_*i*_, *y*_*j*_ *are different from* −, *then y*_*i*_ *< y*_*j*_;
3. *each pair has at least an element that is not* −;
4. *for each j* ∈ [1, *q*] *there is an i such that x*_*i*_ = *j, and for each j* ∈ [1, *l*] *there is an i such that y*_*i*_ = *j*,

Notice that each vertex of *G* is labeled by a single character. The next step is to formally define a recombination.

#### Definition 7

(Recombination). *Let G* = ⟨*V, A, W, λ*⟩ *be a canonical variation graph and let s be an l-long string. Then a recombination is a triple* (*u, v, j*) *where u and v are two vertices of G* = ⟨*V, A, W, λ*⟩, *and j is an integer such that* 1 ≤ *j < l*.

Given a recombination (*u, v, j*) and two walks *w*_1_, *w*_2_ ∈ *W* of *G* = ⟨*V, A, W, λ*⟩, there exists two vertices 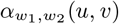 and 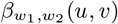 such that (1) *α* strictly precedes *u* in *w*_1_ and *v* in *w*_2_, and for each other vertex 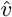 with the same property, 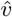 also strictly precedes *α*, and (2) *u* strictly precedes *β* in *w*_1_, *v* strictly precedes *β* in *w*_2_, and for each other vertex 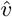 with the same property, *β* also strictly precedes 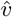. Whenever the two walks *w*_1_ and *w*_2_ are clear from the context, we will omit them.

The intuitive idea is that *α* and *β* are respectively the initial and final vertices of the smallest bubble of *G* = ⟨*V, A, W, λ*⟩ including both *u* and *v*. Such two nodes always exist, since a variation graph has two distinguished source and sink vertices.

#### Definition 8

(displacement). *Let* (*u, v, j*) *be a recombination such that u and v are respectively vertices of the walks w*_1_ *and w*_2_. *Let* 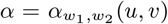*and* 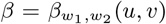. *Let a*_1_ *be the subpath of w*_1_ *from α to u, b*_1_ *be the subpath of w*_1_ *from u to β, a*_2_ *be the subpath of w*_2_ *from α to v, b*_2_ *be the subpath of w*_2_ *from v to β. Then the* displacement *of the recombination is* ||*a*_1_| − |*a*_2_| + 1| + ||*b*_1_| − |*b*_2_| − 1| *and is denoted as* 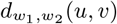.

We will start by introducing the notion of alignment with one recombination, then we will extend the notion to *k* possible recombinations. The main intuition is that a high-quality alignment using a recombination is evidence that the variation graph *G* is lacking a walk that is consistent with the sequence. Using a single recombination means that the sequence and the graph are split into two parts: there is a standard alignment in between the first parts, another standard alignment between the second parts, and the recombination bridges the two parts. We can easily represent this bridge with an (possibly new) arc in the graph. The definition of *α* and *β* allows us to introduce the notion of displacement of a recombination which will be instrumental in computing the penalty associated with a recombination.

#### Definition 9

(Alignment to a variation graph with a recombination). *Let G* = ⟨*V, A, W, λ*⟩ *be a canonical variation graph, and let s be an l-long string. Then an alignment of s to G with a recombination* (*u, v, j*) *is obtained from: (1) two integers r*_1_ *and r*_2_ *with r*_1_ *r*_2_ *such that u is a vertex of* 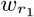 *and v is a vertex of* 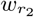; *(2) a subwalk t*_1_ *of w*_1_ *and t*_2_ *of w*_2_, *such that t*_1_ *ends in u and t*_2_ *starts in v*.

*An alignment consists of the concatenation of two alignments against a walk: one between t*_1_ *and s*[: *j*], *and one between t*_2_ *and s*[*j* + 1 :].

The displacement of a recombination models how much the path-preserving alignments are affected by the recombination and it is the difference of the distances of the vertices *u* and *v* with *α* and *β*. Observe that the values +1 and −1, added to |*a*_1_| − |*a*_2_| and to |*b*_1_| − |*b*_2_| in the formula of definition 8 are due to the fact that *v* is the vertex following the vertex *u* in the recombination arc, and a position is added to the right of *u* and subtracted w.r.t. *v*.

#### Example 1.

*Consider the bubble represented in Figure 2. Then the displacement of the recombination is the sum of values* ||*a*_1_| − |*a*_2_| + 1| = |23 + 1| = 0 *and* ||*b*_1_| − |*b*_2_| − 1 = |8 − 8 − 1| = 1, i.e. 1. *An optimal alignment of string s* =*AAGGGGGG to the graph with the recombination results in all matches*.

While any increasing function of the displacement is a reasonable recombination penalty function, we will focus our attention on affine penalties, defined by two parameters *d*_*o*_ (the opening recombination penalty) and *d*_*e*_ (the extending recombination penalty), where the overall penalty is equal to *d*_*o*_ +*d*·*d*_*e*_, where *d* is the displacement of the recombination. Now we can extend Definition 9 to allow *k* recombinations.

#### Definition 10

(Alignment with *k*-recombinations). *Let G* = ⟨*V, A, W, λ*⟩ *be a canonical variation graph and let s be an l-long string. Then an alignment of s to G with k recombinations* (*u*_*i*_, *v*_*i*_, *j*_*i*_) *with* 1 ≤ *j*_*i*_ *< j*_*i*+1_ *< l consists of the concatenation of the k* + 1 *alignments between s*[*ji*−1 + 1 : *ji*] *and the subwalk t*_*i*_, *such that: (1)* ⟨*t*_1_, …, *t*_*k*+1_⟩ *is a sequence of walks of G, where each t*_*i*_ *is a subwalk of* 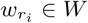, *such that: (1) r*_*i*_ *is a sequence of k* + 1 *integers such that r*_*i*_ *r*_*i*+1_, *u*_*i*_ *is a vertex of* 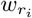*and v*_*i*_ *is a vertex of* 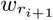, *for each* 1 ≤ *i* ≤ *k, and*

a. *the subwalk t*_1_ *starts with the source of G and t*_*k*+1_ *ends with the sink of G;*
b. *all vertices α*(*u*_*i*_, *v*_*i*_) *and all vertices β*_*i*_(*u*_*i*_, *v*_*i*_) *are distinct*.
c. *for each i, the subwalk t*_*i*_ *ends with the vertex v*_*i*_ *and t*_*i*+1_ *starts with the vertex u*_*i*+1_. *The value of such alignment incorporates also the penalties of all k recombinations*.

The main idea is that *k* recombinations corresponds to *k* + 1 subwalks and *k* recombination arcs connecting the end of a subwalk and the beginning of the subsequent subwalk. Moreover, the entire alignment and the sequence are both split into *k* + 1 portions, so that the *i*-th portion of the sequence is aligned (without recombinations) against the *i*-th subwalk, and such alignment is the *i*-th portion of the alignment. The natural computational problem is to compute an alignment with at most *k* recombination and with maximum value, where the objective function is the score of the alignment minus the gap penalties, minus the recombination penalties.

### 3.2 Aligning against a variation graph

First we will consider the case where we want an optimal alignment without recombinations. A trivial approach is to extract the sequences corresponding to the walks and align the input sequence against each of those with the Needleman-Wunsch [17] or the more recent wavefront alignment [15] algorithms. Anyway, this approach does not exploit the fact that the pangenome is stored as a graph. An alternative algorithm is a variation of the approach taken by POA [12] on an acyclic variation graph *G* = ⟨*V, A, W, λ*⟩. By Lemma 1, we can assume that *G* is canonical.

In the following, let *M* [*v, i, p*] be the optimal score of the global alignment between the initial portion of the path *p* ∈ *W* that ends in the vertex *v* and the *i*-long prefix *r*[: *i*] of the read *r*. We can describe the values of the matrix *M* with the usual recurrence equation where, for simplicity, we denote with *g* the penalty of an indel, with *m* the score of a match, and with 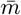 the score of a mismatch:

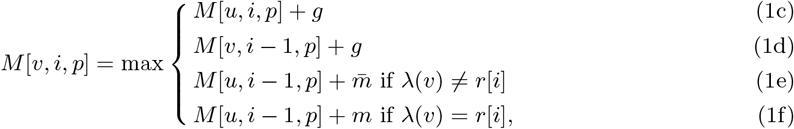

where *λ*(*v*) is the character labeling the vertex *v* and *u* is the vertex preceding *v* in *p*. Moreover, *M* [*s*, 0,·] = 0 if *s* is the source of *G*. Since there can be several paths in *W* traversing the same vertex, we can avoid some redundant computation by considering them together. More precisely, for each vertex *v* we choose a path traversing *v*: such a path is called the *reference path* and is denoted by *α*(*v*).

We introduce another matrix *D*[*v, i, p*] defined as *D*[*v, i, p*] = *M* [*v, i, p*] − *M* [*v, i, α*(*v*)], that is the matrix *D* stores differences of values of *M* with respect to a reference path. Observe that *D*[*v, i, p*] may be also a negative value. Since we expect the values of the matrix *D* to be small, we can encode them compactly (e.g. with an Elias gamma encoding), reducing the memory occupation. By construction, *M* [*v, i, p*] = *D*[*v, i, p*] + *M* [*v, i, α*(*v*)], hence we can store only the values *D*[*v, i, p*] and not the values *M* [*v, i, p*] for such pairs (*p, v*). The following proposition, relates two paths traversing the same arc.

#### Proposition 1

(distance preserving). *Let p*_1_ *and p*_2_ *be two paths traversing the arc* (*v, w*), *and let r*[: *j*] *be a prefix of the string. Assume that the cases 1e, 1f that achieve the maximum for M* [*w, j, p*_1_] *are exactly those achieving the maximum for M* [*w, j, p*_2_]. *Then M* [*w, j, p*_1_] − *M* [*w, j, p*_2_] = *M* [*v, j* − 1, *p*_1_] − *M* [*v, j* − 1, *p*_2_].

A consequence of Proposition 1 is that we do not need to recompute some values of the matrix *D*. In fact, given an arc (*v, w*) such that *α*(*v*) = *α*(*w*) and computing the value of *M* [*w, i, α*(*v*)] does not introduce gaps, then *D*[*w, i, p*] = *D*[*v, i* − 1, *p*]. Another optimization stems from an iterative application of the above observation. This fact can be exploited to speed up the computation on non-branching paths of the graph, if no indel is introduced in aligning that portion of the path and the corresponding portion of the sequence. More precisely, if a path *w*_1_, …, *w*_*q*_ of the graph consists of vertices *w*_*i*_ with exactly one incoming and one outgoing arcs, then *D*[*w*_1_, *j, p*] = *D*[*w*_*i*+1_, *j* + *i* − 1, *p*] for all *i* ≥ 1. Therefore, we only need to store the values *D*[*w*_1_, *j, p*].

Notice that the optimizations that we have described hold for any choice of the reference paths. We will describe how we actually compute such paths and, most importantly, how to quickly compute the relevant values of the matrix *D* — remind that equation 3.2 holds only when the reference is the same in both vertices *v* and *w* for a given arc (*v, w*). Let us consider a vertex *w*, the set of paths traversing *w* and the set *N* of vertices of that are the initial endpoint of an arc ending in *w* (that is, *N* = {*v* : (*v, w*) ∈ *E* ∧ *v* ∈}). Equation 3.2 suffices when |*N* | = 1 and *α*(*v*) ∈ : in this case, simply pose *α*(*w*) = *α*(*v*). Essentially there are two more cases to consider: at least one reference path *p* in a vertex of *N* traverses *w*, or no such reference path exists. In the first case, pick any such path *p* and pose *α*(*w*) = *p*. Moreover let *v* be the vertex of *N* such that *v* ∈ *N*. Equation 3.2 still applies to all paths *p* that traverse the arc (*v, w*). For all other paths *p, D*[*w, i, p*] = *M* [*w, i, p*] − *M* [*w, i, α*(*w*)]. In the second case, that is when no suitable reference path exists, we pick any path *p* ∈ and pose *α*(*w*) = *p*. Just as for the first case, *D*[*w, i, p*] = *M* [*w, i, p*] − *M* [*w, i, α*(*w*)] for all path *p* ≠ *α*(*w*).

We describe a dynamic programming approach for computing optimal alignments with at most one recombination. We recall that the instance of such problem consists of a variation graph *G* = ⟨*V, E, W* ⟩, a *n*-long string *s*, a score matrix *d*, a gap penalty (*g*_*o*_, *g*_*e*_), and a recombination penalty (*d*_*o*_, *d*_*e*_). An optimal alignment can have zero or one recombination. In the first case, we have already described the recurrence equation describing the optimal solution. In the second case, we have an optimal path-preserving alignment, without any recombination, between a prefix of the sequence and an initial portion of a path, and a second path-preserving optimal alignment, without any recombination, between a suffix of the sequence and a final portion of a path. Those two alignments are connected by a recombination.

We know how to compute the matrix *M* [*v, i, p*], *i*.*e*. the optimal score of the global alignment between the initial portion of the path *p* ∈ *W* that ends in the vertex *v* and the *i*-long prefix *s*[: *i*] of the sequence *r*. A similar recurrence equation describes the matrix *R*[*v, i, p*] that is the optimal score of the global alignment between the final portion of the path *p* ∈ *W* that starts in the vertex *v* and the suffix *s*[*i* :] of the sequence *r* starting in position *i*. The two matrices can be computed in parallel. The value of an optimal alignment with at most one recombination between *s* and *G* is given by the following equation where *s*_*i*_ and *s*_*e*_ respectively are the source and the sink of *G* and *n* is the length of the string *s*.

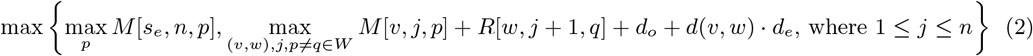

If the maximum is achieved by the first maximum, then the optimal alignment is obtained without any recombination. Otherwise there exists two paths *p, q* ∈ *W*, and two nodes *v, w* ∈ *V* such that the best alignment of the sequence *s* against the graph *G* is given by the subpath of *p* from node *s*_*i*_ to *v*, and the subpath of *q* from node *w* to *s*_*e*_ (plus the affine displacement penalty), meaning that the recombination corresponds to the arc (*v, w*). The running time of the naïve algorithm exploiting 2 is *O*(|*V* |^2^|*W* |^2^*n*).

## 4 Experimental results

We implemented the method described in Section 3.2 in our tool `RecGraph` (available at https://github.com/AlgoLab/RecGraph under the MIT license) which computing optimal alignments against a sequence graph (called unrestricted mode), against variation graphs, but without recombinations, (path-preserving mode), and against variation graphs, but with a recombination (recombination mode).

The focus of our experimental evaluation (https://github.com/AlgoLab/RecGraph-exps) is to assess the effectiveness of introducing recombinations in obtaining better alignments. To this aim, we considered *Escherichia coli*, a model bacterium for which the study of the pangenome has proven useful [8, 3], and we simulated the scenario where reads from a novel recombinant strain have to be aligned to the pangenome of already-known strains. We randomly selected 50 *E. coli* core genes from the `panX` platform [8] and we created 50 pangenome graphs using the `make prg` utility from `Pandora` [3]. Since `make prg` produces sequence graphs (i.e., GFA files with no path lines), we added such information by aligning back each input strain to the corresponding pangenome graph using `RecGraph` in the unrestricted global mode and by considering the computed alignments as paths of the graph. In such a way, we have been able to obtain a variation graph for each considered gene. Starting from these graphs, we split the strains (paths) of each gene (graph) in two sets: the set of known strains (i.e., paths that will be kept in the final variation graph) and the set of new recombinant strains (i.e., paths that will be removed from the graph and used to simulate reads). We recall that our goal was to align reads from a possibly new recombinant strain to the set of known strains. To avoid modifying the set of edges, for each graph, we greedily computed a minimal subsets of paths covering all the edges and we considered them as the *known strains*, whereas all the remaining paths as *recombinant strains*. Each recombinant strain is seen as a mosaic of the known strains. Out of the 50 considered genes, we obtained 616 recombinant strains. From each recombinant strain, we simulated 19 124 15x Illumina paired-end reads (with read length 150bp) using `dwgsim` (http://github.com/nh13/dwgsim). To compute our baseline, we aligned each sample against the corresponding recombinant strain using `BWA mem` [13]. We consider as baseline the Levenshtein distance between the read and the substring of the recombinant strain which `BWA` maps the read to. With a slight abuse of language, we call such a distance as *edit distance of the alignment*.

We aligned each sample using `RecGraph` to the minimal variation graph (i.e., the graph where the paths are only the known strains) in path-preserving mode and in recombination mode. To put our results in perspective, we also aligned each sample to the minimal variation graph using `giraffe` [21]. Since `RecGraph` in path-preserving mode aligns to a variation graph without allowing recombinations, its alignments should be similar to those obtained with `giraffe`. On the other hand, since `RecGraph` in recombination mode can introduce recombinations in the alignments, its alignments should be more similar to those produced by `BWA`, that aligns reads directly to the corresponding recombinant strain.

We evaluated the accuracy of the tools in terms of edit distance of the alignments they produce compared with that of the alignments computed by `BWA`. Since `giraffe` has not been able to correctly align both mates of 7 pairs, we decided to remove such pairs from the analysis. This resulted in 19 117 analyzed pairs. Table 1 summarizes the comparison when alignments cannot introduce recombinations. In this case, the edit distance of the alignments computed by `RecGraph` in path-preserving mode for 17 281 read pairs (out of 19 117) matches that of the alignments computed by `BWA`. Only 390 alignments (2%) have edit distance smaller than that of `BWA`. Manual analysis of some of these alignments revealed that the presence of sequencing errors that match the SNP alleles present in other known strains led the tools to align the reads to a different strain. This is expected as the reads are short, hence they overlap to only a small number of SNPs. Interestingly, this fact further confirms the advantage of using graph representations of the pangenome, as these reads likely maps to vertices shared by several paths. Hence, a post-processing of the alignments can easily detect the error by computing the observed coverage of each vertex. Instead, 1 446 alignments (7.5%) have edit distance larger than that of `BWA`. This is the case of reads overlapping the recombination. Those reads cannot be aligned to the recombinant strain in path-preserving mode since that strain is not one of the distinguished paths of the variation graph. The edit distance of the other alignments matches that of `BWA`, likely indicating that reads are mapped to the true location and that they do not overlap any recombination. `giraffe` behaves similarly to `RecGraph`, with small differences probably due to the heuristic nature of the method (`RecGraph` is exact).

**Table 1:** Number of read pairs whose alignment to the graph computed by `giraffe` and `RecGraph` in path-preserving mode have edit distance less than/equal to/greater than that of the alignment computed by `BWA` w.r.t. the corresponding recombinant strain.

|  | **giraffe** | **RecGraph: path-preserving** |
| --- | --- | --- |
| Less than BWA | 390 | 390 |
| Equal to BWA | 17 272 | 17 281 |
| Greater than BWA | 1 455 | 1 446 |

The assessment of the path-preserving alignments empirically highlights that this kind of alignments are sub-optimal in a small, but significant, portion of the sample, hence supporting the need to explicitly model recombinations in sequence-to-graph alignments. As a consequence, we then evaluated the impact of introducing recombinations in the alignments. We chose to fix the opening recombination penalty (*d*_*o*_) to 4 and the extending recombination penalty (*d*_*e*_) to 0.1. Such a choice of values, along with a mismatch penalty of 4 and a match score of 2, implies that a recombination is less costly than a mismatch if the displacement is at most 20. We argue that this choice is adequate since the samples we considered are composed of (simulated) Illumina short reads. For the same reason, and because a recombination can be cheaper than a mismatch, we chose to avoid introducing recombinations in the 15bp-long prefix or suffix of the read. Indeed, we argue that alignments that include a recombination near the begin or the end of the read are usually incorrect, since sequencing errors can be easily confounded with SNPs close to read extremities, therefore the support for such a recombination is insufficient. Table 2 summarizes how read alignments changed their edit distance from path-preserving mode to recombination mode. In particular, we computed the number of alignments whose edit distance *in path-preserving mode* is smaller than/equal to/greater than that of the alignments computed by `BWA` and the edit distance of the alignment *in recombination mode* is smaller than/equal to/greater than that of the alignments computed by `BWA`. The first observation is that only 75 pairs have an alignment with (possibly) a recombination whose edit distance is greater than that of the alignment computed by `BWA`, out of the 1 446 read alignments in path-preserving mode. Part of these 75 pairs have “suboptimal” alignments in both path-preserving and recombination mode since the recombination that allows to improve the edit distance is located near one of the ends of the read and that was filtered out as explained above. Notably, the vast majority (1 309 out of 1 446) suboptimal alignments in path-preserving mode become optimal (in the sense that they match the edit distance computed by `BWA`) in recombination mode. This suggest that, by introducing a recombination, the alignment matches that of the recombinant strain used as reference in `BWA`. Finally, 62 suboptimal alignments in path-preserving mode reduce their edit distance under that of `BWA`. This is likely due to sequencing errors that fortuitously match SNP alleles of other strains. As argued before, a simple post-processing of the computed alignments should easily identify (and filter out) these spurious matches based on the observed vertex coverages. On the other hand, 1 036 pairs whose edit distance of the alignments in path-preserving mode was equal to that of the alignments of `BWA` have an edit distance in recombination mode lower than that of `BWA`. Unfortunately, this is a inherent limit of using short-read sequencing technologies, as they can only provide linkage evidence for short spans of the pangenome.

**Table 2:**
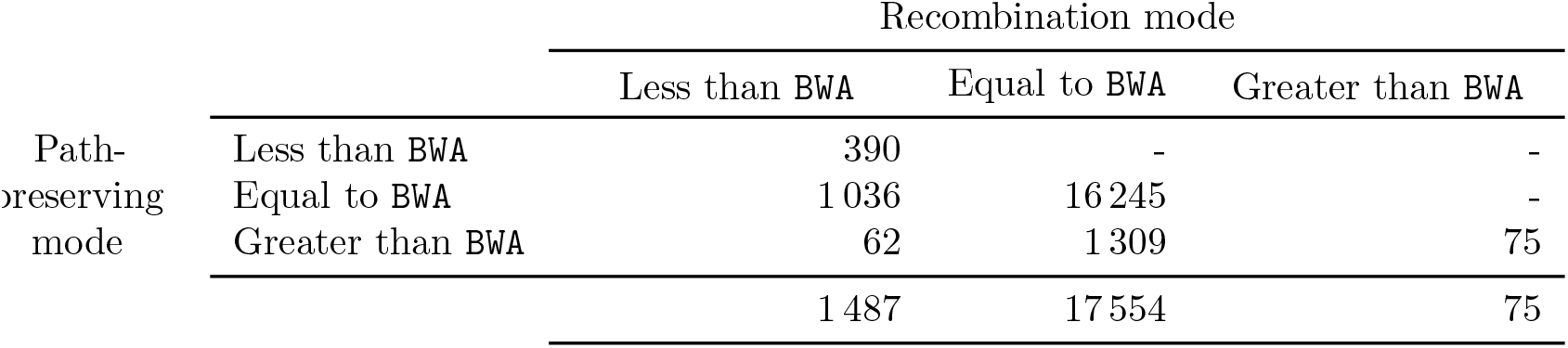
Impact of allowing a recombination in the alignment. Each cell contains the number of read pairs whose alignment in *path-preserving* mode have edit distance less than/equal to/greater than that of the alignment computed by `BWA` (row headers) and whose alignment in *recombination* mode have edit distance less than/equal to/greater than that of the alignment computed by `BWA` (column headers). For example, there are 1 309 read pairs whose alignment in path-preserving mode had edit distance greater than that of `BWA` but whose alignment in recombination mode has the same edit distance of the alignment computed by `BWA`.

|  |  | Recombination mode |  |  |
| --- | --- | --- | --- | --- |
|  |  | Less than BWA | Equal to BWA | Greater than BWA |
| Path-preserving mode | Less than BWA | 390 | - | - |
|  | Equal to BWA | 1036 | 16 245 | - |
|  | Greater than BWA | 62 | 1 309 | 75 |
|  |  | 1 487 | 17 554 | 75 |

All experiments on a 64bit Linux (Kernel 5.15.0) system equipped with two AMD® Epyc 7301 processors and 128 GB of RAM. `RecGraph` in recombination mode took from few seconds to 15 minutes depending on the input graph size. We remind that our approach guarantees to find an optimal solution and that there are several heuristics that can be applied to speed up the computation — eventually forgoing the guarantee in a few cases. As expected, `RecGraph` in recombination mode is more time consuming than both its path-preserving counterpart and `giraffe`, which both took from few seconds to half a minute. All tested tools required less than 512MB of memory.

## 5 Conclusions and open problems

We have started an investigation of incorporating additional events, such as a recombination, in a sequence-to-graph alignments. This new notions of alignments can be crucial to investigate the presence of biological events that contribute to the exchange of genetic material among individuals of the same species such as homologous recombinations or horizontal gene transfer. This phenomenon is mainly of interest in bacterial genomes that are characterized by a higher degree of recombination events.

We have formalized the notion of alignment with recombination in a variation graph, designed a dynamic programming algorithm for the problem, and have implemented it in RecGraph. An experimental study over a bacterial pangenome shows that RecGraph is effective in obtaining high-quality alignments of sequences that can be expressed only as a mosaic recombination of paths of the graph/ The alignment with recombinations poses new theoretical challenges in the general problem of mapping sequences to a graph. For example, an open problem is the efficient computation of the displacement of all possible recombinations, as that would improve time complexity of computing equation 2. A further direction is to use RecGraph as a building block for developing more efficient heuristic aligners with recombinations.

## Appendix

We introduce a formal notion of graph equivalence, based on the idea that two equivalent graphs must have walks encoding the same genomes. Based on this notion, we can show that there exists a canonical variation graph *G*_*c*_ equivalent to any given variation graph *G*, as stated in Lemma 1.

### Definition 11

(Equivalent variation graphs). *Given G*_1_ = ⟨*V*_1_, *A*_1_, *W*_1_, *λ*_1_⟩ *with λ*_1_ : *V*_1_ → Σ^+^, *and G*_2_ = ⟨*V*_2_, *A*_2_, *W*_2_, *λ*_2_⟩ *with λ*_2_ : *V*_2_ → Σ^+^ *two variation graphs, then G*_1_ = ⟨*V*_1_, *A*_1_, *W*_1_, *λ*_1_⟩ *and G*_2_ = ⟨*V*_2_, *A*_2_, *W*_2_, *λ*_2_⟩ *are* equivalent if the two multisets {*λ*(*w*) : *w* ∈ *W*_1_} and {*λ*(*w*) : *w* ∈ *W*_2_} are the same.

### Lemma 1.

*Let G* = ⟨*V, A, W, λ*⟩ *be an acyclic variation graph. Then there exists a canonical variation graph G*_1_ = ⟨*V*_1_, *A*_1_, *W*_1_, *λ*_1_⟩ *that is equivalent to G*.

