## Supplementary material for "RecGraph: adding recombinations to sequence-to-graph alignments"

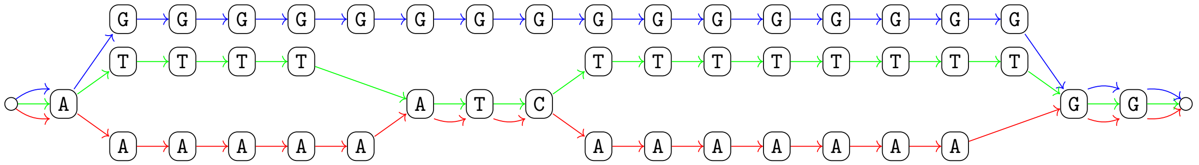

**Figure 4:** Example of canonical variation graph with three paths (one for each color).

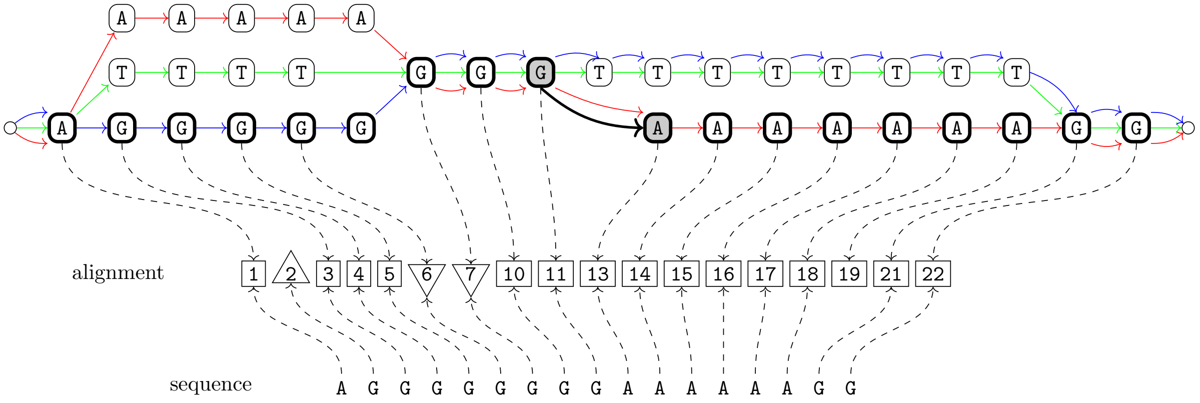


**Figure 5:** Example of path-preserving alignment of a sequence against a variation graph with a recombination. The recombination is represented by the thick black arc connecting two vertices with gray background. This example shows a recombination arc that is parallel to an already existing arc.

### Appendix

We introduce a formal notion of graph equivalence, based on the idea that two equivalent graphs must have walks encoding the same genomes. Based on this notion, we can show that there exists a canonical variation graph $G_c$ equivalent to any given variation graph $G$, as stated in Lemma 1.

**Definition 11** (Equivalent variation graphs)**.** *Given* $G_1 = \langle V_1, A_1, W_1, \lambda_1 \rangle$ *with* $\lambda_1 : V_1 \to \Sigma^+$*, and* $G_2 = \langle V_2, A_2, W_2, \lambda_2 \rangle$ *with* $\lambda_2 : V_2 \to \Sigma^+$ *two variation graphs, then* $G_1 = \langle V_1, A_1, W_1, \lambda_1 \rangle$ *and* $G_2 = \langle V_2, A_2, W_2, \lambda_2 \rangle$ *are* equivalent if the two multisets $\{\lambda(w) : w \in W_1\}$ and $\{\lambda(w) : w \in W_2\}$ are the same.

**Lemma 1.** *Let* $G = \langle V, A, W, \lambda \rangle$ *be an acyclic variation graph. Then there exists a canonical variation graph* $G_1 = \langle V_1, A_1, W_1, \lambda_1 \rangle$ *that is equivalent to* $G$*.*
